# Cerebellar granule neurons induce Cyclin D1 in an early stage of Huntington’s disease

**DOI:** 10.1101/2022.11.08.515613

**Authors:** Susanne Bauer, Chwen-Yu Chen, Maria Jonson, Lech Kaczmarczyk, Srivathsa Magadi, Walker S. Jackson

## Abstract

Although Huntington’s disease (HD) is predominantly defined by selective vulnerability of striatal projection neurons, there is increasing evidence that cerebellar degeneration modulates clinical symptoms. However, little is known about cell type-specific responses of cerebellar neurons in HD. To dissect early disease mechanisms in the cerebellum and cerebrum, we analyzed translatomes of neuronal cell types from both regions in a new HD mouse model. For this, HdhQ200 knock-in mice were backcrossed with the calm, 129S4 strain, to constrain experimental noise caused by variable hyperactivity of mice in a C57BL/6 background. Behavioral and neuropathological characterization showed that these S4-HdhQ200 mice had very mild behavioral abnormalities starting around 12 months of age that remained mild up to 18 months. By 9 months, we observed abundant Huntingtin-positive neuronal intranuclear inclusions (NIIs) in the striatum and cerebellum. The translatome analysis of GABAergic cells of the cerebrum further confirmed changes typical of HD-induced striatal pathology. Surprisingly, we observed the strongest response with 626 differentially expressed genes in glutamatergic neurons of the cerebellum, a population consisting predominantly of granule cells, a cell type commonly considered disease-resistant. Our findings suggest vesicular fusion and exocytosis, as well as differentiation-related pathways are affected. Furthermore, increased expression of cyclin D1 (*Ccnd1*) in the granular layer and upregulated expression of polycomb group complex protein genes and cell cycle regulators *Cbx2, Cbx4* and *Cbx8* point to a putative role of aberrant cell cycle regulation in cerebellar granule cells in early disease.

## INTRODUCTION

Huntington’s disease (HD) [OMIM:143100] is a progressive and fatal neurodegenerative disease caused by a CAG repeat expansion in exon 1 of the Huntingtin gene (*HTT*), which results in an expanded polyglutamine stretch in the Huntingtin protein (HTT). While both wild type and mutant HTT (mHTT) are expressed ubiquitously, the striatum is the region most affected in HD. In particular, GABAergic striatal projection neurons (SPNs) show pronounced selective vulnerability, with early degeneration of dopamine 2 receptor (Drd2)-expressing SPNs of the indirect pathway, followed by Drd1-expressing SPNs of the direct pathway (Reiner et al., 1988). In addition to striatal degeneration, renewed attention has recently been placed on cerebellar pathology and reassessment of its potential role in clinical symptoms of HD. The cerebellum plays a role in functions commonly affected in HD patients, such as impairment of dexterity, postural instability, or ataxia-like symptoms. Indeed, the presence of neuropathology in the cerebellum has been described in HD patients and mouse models, predominantly of Purkinje cell loss, while the granular layer was relatively spared (Jeste et al., 1984; Dougherty et al., 2013; Rüb et al., 2013; Singh-Bains et al., 2019). Neuroimaging showed degeneration of cerebellar gray matter in anterior and posterior lobules of the cerebellum in patients with mild motor symptoms (Padron-Rivera et al., 2021). Similarly, significant loss of Purkinje neurons was found in patients with motor phenotypes specifically, corroborating the involvement of the cerebellum in HD symptoms and a correlation of HD phenotype with cerebellar pathology (Singh-Bains et al., 2019). Interestingly, there is increased functional connectivity between the striatum and cerebellum in asymptomatic children carrying CAG expansions, which declines in an age and CAG repeat length-dependent manner (Tereshchenko et al., 2020). Thus, the cerebellum may help to maintain normal motor function in early HD, compensating for the loss of striatal function (Van Der Plas et al., 2020). Consequently, the spectrum of symptoms in HD may be influenced by loss of cerebellar compensation in addition to striatal dysfunction, making the cerebellum an interesting therapeutic target in HD.

Despite the importance of cerebellar disturbances in HD, the mechanisms behind it remain largely unclear. For example, the occurrence and degree of cerebellar pathology does not strictly correlate with CAG repeat length or the degree of striatal degeneration (Rüb et al., 2013; Singh-Bains et al., 2019; Franklin et al., 2021), which begs the question what other mechanisms drive cerebellar degeneration in HD. Recent studies highlighted early gene expression responses in striatal cell types (Lee et al., 2020), however, similar studies of the cerebellum are missing. We therefore studied cell type-specific responses of excitatory (vGluT2^+^) and inhibitory (Gad2^+^) cerebellar populations in an early, pre-symptomatic disease stage, using cell type-specific translatome analysis with RiboTag (Sanz et al., 2009) in our S4-HdhQ200 model. Surprisingly, cerebellar vGluT2^+^ neurons showed the strongest response with 626 differentially expressed genes (DEGs) which were enriched for vesicular exocytosis and included upregulation cell cycle regulators Cyclin D1 and chromobox (Cbx) protein genes. This study suggests granule cells, commonly considered resistant in HD, may be affected early in disease.

## RESULTS

Although conventional transgenic mice can cause severe and early disease onset, accelerating experimentation, such models are vulnerable to expression pattern artifacts (Kaczmarczyk and Jackson, 2015). We therefore employed a knock-in mouse model. The mouse model carrying a repeat of approximately 200 CAGs in the endogenous gene (HdhQ200) was initially established in a C57BL/6J background (Lin et al., 2001; Heng et al., 2010). Since approximately 20% of mice in this background demonstrate nocturnal hyperactivity (Kaczmarczyk et al., 2021), which could create gene expression noise, we backcrossed the mutant allele into the 129S4 background. To characterize the disease phenotype in this new genetic background, we performed behavioral and histological analyses on S4-HdhQ200 heterozygous mice and littermate controls (hereafter HD and control respectively) at different ages (**Fig. S1**).

### HD mice in the S4 background are mildly affected

To assess behavioral changes during disease progression, we evaluated animals between the ages of 3-18 months using standard paradigms. At each experimental time point, mice performed a session on the accelerating rotarod, followed by walking on a balance beam, 1 h of burrowing and a second balance beam task. Additionally, body weight was monitored over the course of the behavioral assessment. Neither males nor females showed a significant difference in body weight between HD and controls (**Fig. 1 A, B**), although female HD mice tended to weigh less. Rotarod performance declined with aging for both control and HD mice, but significant differences were only observed at 15 months (p < 0.05; **Fig. 1 C**). Some HD mice performed very poorly on the balance beam task beginning at 12 months, but the majority performed as well as controls and the group average was similar to controls (**Fig. 1 D, F**). Burrowing is an innate, instinctive, and rewarding behavior in mice, and burrowing paradigms are used as an assessment of overall well-being and effects of disease on the performance of spontaneous behavior (Deacon, 2009). HD mice burrowed less at 9 and 12 months (p < 0.05). This trend continued at 15 and 18 months, but since several HD mice performed as well as controls, the differences were not significant (**Fig. 1 E**). A larger cohort of mice may have produced more statistically significant differences, but it would not have changed the overall conclusion that S4-HdhQ200 mice showed only a very mild behavioral phenotype, even at an advanced age, similar to when this mutation is in the C57Bl/6 background (Lin et al., 2001; Heng et al., 2010).

**Figure 1:**
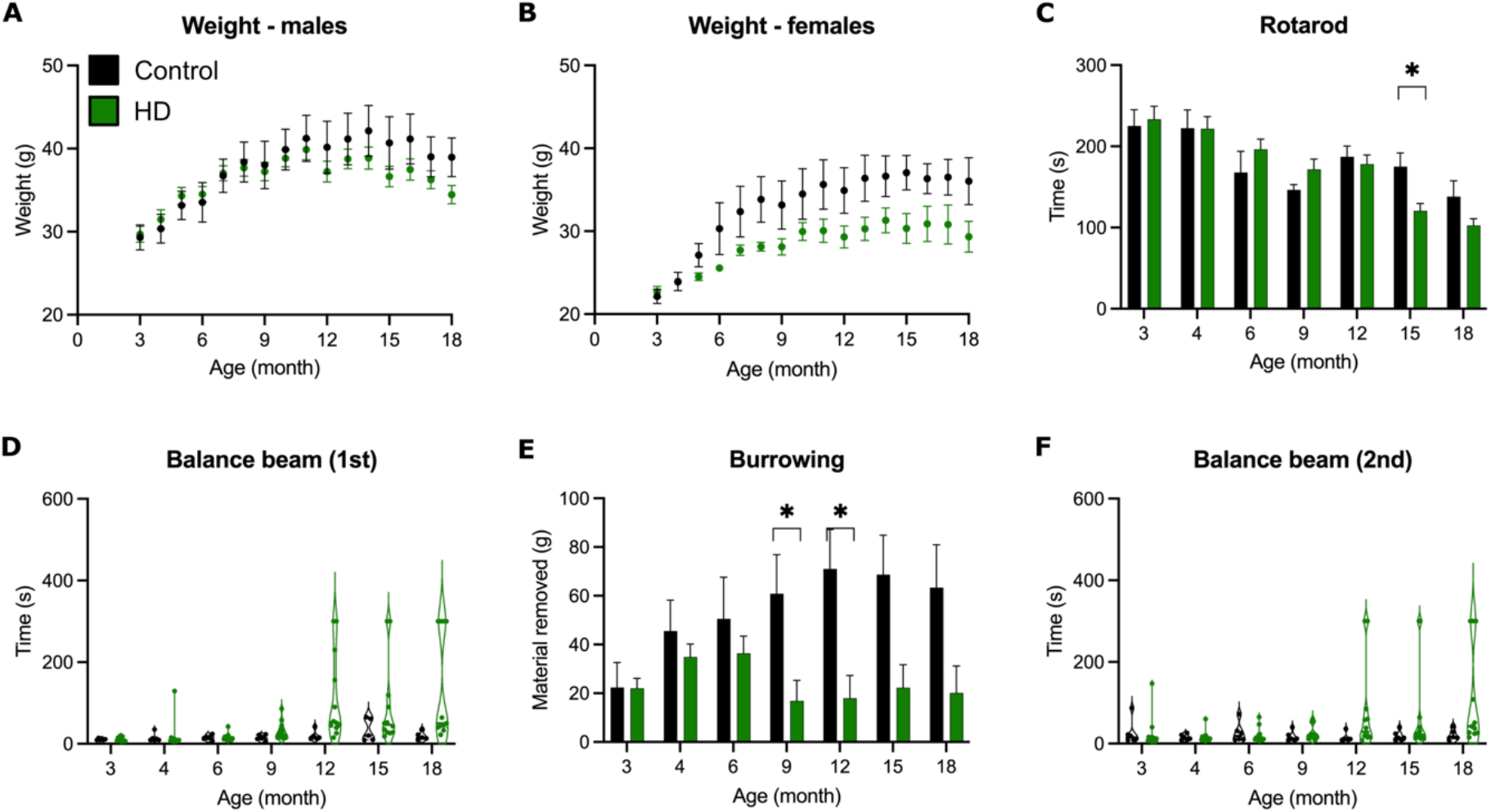
S4-HdhQ200 mice have a mild behavioral phenotype. **(A, B)** Body weight is not significantly reduced in male (A), or female (B) HD mice compared to littermate controls. **(C)** HD mice remain on the rotarod for less time than control mice at 15 months. **(D, F)** HD mice show no significantly different performance in balance beam task. For each time point, two repeat measures were performed, with a one-hour burrowing period between the first **(D)** and second **(F)** test. **(E)** HD mice burrow less than littermate controls at 9 months and 12 months. HD n = 12 (green), control n = 7 (black), unpaired t-test without corrections for multiple testing, * represents p < 0.05.

In contrast to the mild behavioral phenotype, the histopathological phenotype revealed a clear neurodegenerative process. NIIs are common in polyglutamine diseases, such as HD and several spinocerebellar ataxias, and their extent can be indicative of disease stage. To improve detection of Htt protein aggregates, soluble Htt was partially removed by applying a mild proteinase K treatment, resulting in bright puncta on a background of diffuse staining, only in HD mice (**Fig. 2 A, B**). The proportion of cells with proteinase K-resistant Htt aggregates was measured at 4, 9 and 18 months in eight brain regions (**Fig. 2 C**). By 9 months of age aggregates were detected in all brain regions, however hippocampus, striatum, and cerebellum had the highest percentage of cells with aggregates (**Fig. 2 D, Fig. S2**). Immunofluorescent co-staining of Htt with the nuclear stain DAPI revealed that nearly all aggregates in these brain regions were NIIs (**Fig. 2 E, Fig S2**). In contrast, aggregates in the thalamus, brain stem, midbrain, and hypothalamus were much less frequent and presented mostly as cytosolic aggregation foci, appearing like NIIs but not within a nucleus (**Fig. 2 D, E; Fig S2**). To determine if regions with high NII load expressed *Htt* the highest, *in situ* hybridization (ISH) for *Htt* mRNA was performed. Surprisingly, the striatum and cerebellum were among the lowest expressing regions, whereas the thalamus was the second highest expressing region, indicating something other than expression levels determines NII load (**Fig 2F**). We also observed a significant, age-dependent downregulation of *Htt* mRNA levels in HD mice in all brain regions except the hypothalamus by 9 months (**Fig. 2 F; Fig. S2**).

**Figure 2:**
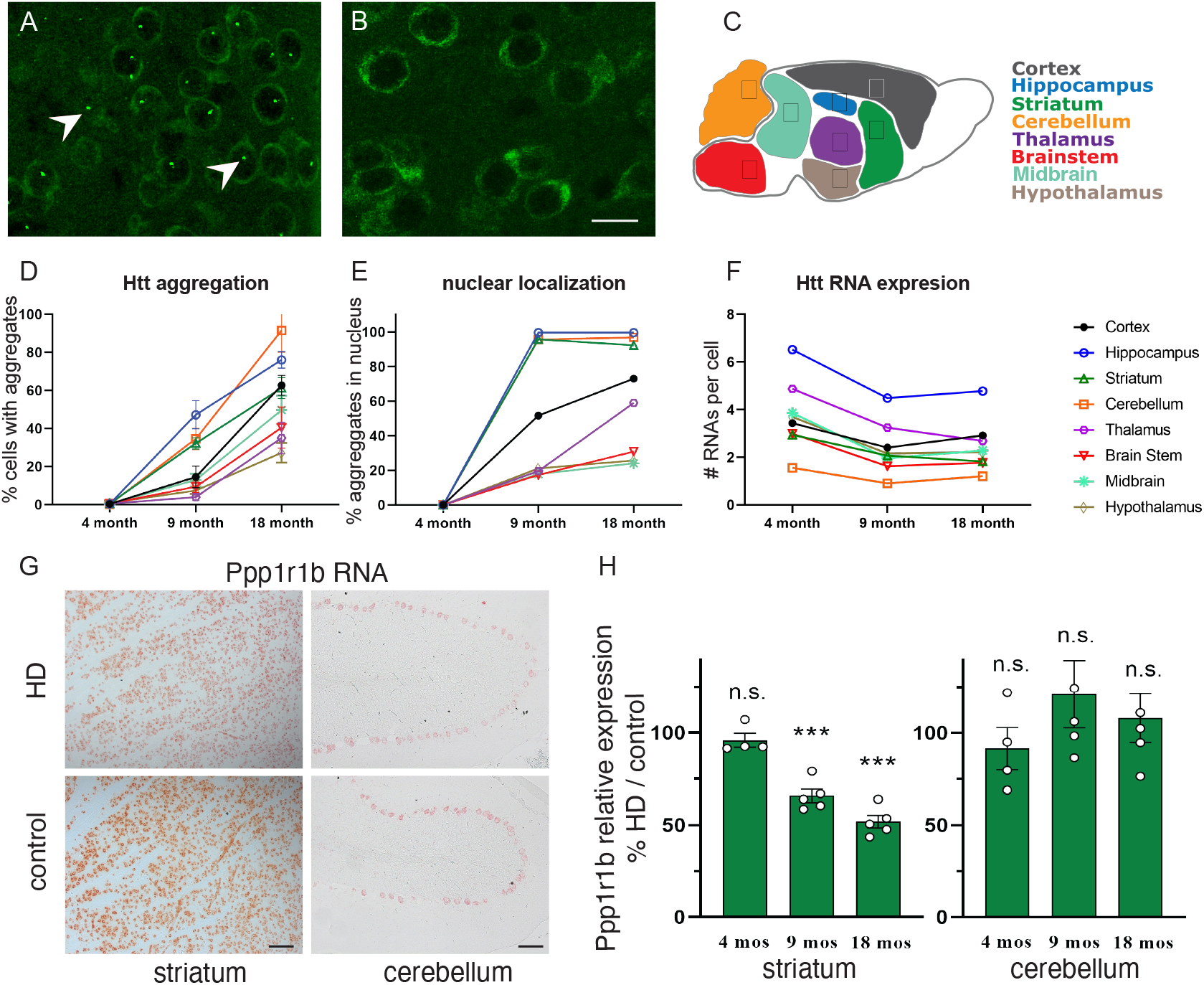
S4-HdhQ200 show HD-typical neuropathology by 9 months of age. **A)** Htt aggregates (2 examples marked with arrowheads) present in 9 months (mos) old HD striatum **(A)**, were absent from 9 mos old control striatum **(B). C)** Overview of brain regions analyzed. **D-F)** Proportion of aggregates per cell **(D)**, percent of aggregates that are NIIs **(E)**, and the number of RNA molecules detected per cell **(F)**. All charts share the key in F, using the same color scheme in **C**; statistical analyses are presented in figure S2. **G-H)** Representative images of RNA ISH for Ppp1r1b in 9 mos old brain sections **(G)**, and quantification and analysis in **H**. Unpaired t-test without corrections for multiple testing, *** represents p < 0.001. Scale bar in B represents 25 μm, the scale bars in G represent 100 μm.

Previous analyses of similar HD knock-in mouse models revealed downregulation of common HD-associated genes in striatal cell types early in the disease (Langfelder et al., 2016; Lee et al., 2020), suggesting loss of striatal identity may be one of the earliest features of HD pathology. We therefore performed ISH staining for one such marker, Ppp1r1b (DARPP32), which is abundant throughout the striatum and in Purkinje neurons of the cerebellum. Ppp1r1b was reduced 38% in the striatum at 9 months, and 51% by 18 months but was unchanged in the cerebellum at all time points (**Fig. 2 G, H**). Staining for Gfap and Iba1 showed no obvious signs of astrogliosis or microgliosis at any disease stage (**Fig. S3**). Finally, no differences were detected between HD and control mice at 4 months of age, indicating the abnormalities detected at 9- and 18-months resulted from a degenerative process. In summary, like other HD knock-in mice, S4-HdhQ200 mice have very mild behavioral and neuropathological changes, with abundant NII accumulation in specific brain regions and Ppp1r1b reduction in the striatum at 9 months. Therefore, here we considered 9 months as an ideal early disease stage for assessment of cell type-specific translatome changes.

### Cell type-specific translatome analysis at 9 months

To study cell type-specific translatomes we employed the RiboTag method (Sanz et al., 2009). It utilizes a knock-in mouse line in which the endogenous gene encoding large subunit ribosomal protein 22 (Rpl22) has been engineered such that activation by Cre recombinase results in an HA antibody epitope being encoded on the C-terminus of Rpl22, and consequently HA-tagged ribosomes (RiboTag). From such samples one can immunoprecipitate (IP) HA-tagged ribosomes from cell types of interest, and sequence the attached translating mRNAs, representing the translatome. To this end we crossed Hdh^Q7/Q200^-Rpl22HA^flox/flox^ mice with homozygous Cre-driver lines to generate HD and control mice expressing RiboTag in targeted cell types. We targeted general populations of either GABAergic (Gad2^+^) or glutamatergic (vGluT2^+^) neurons in the cerebellum and the cerebrum, where cerebrum is the remaining part after removal of the cerebellum and olfactory bulb from the brain. In the cerebrum, we also targeted the subset of GABAergic neurons expressing the neuropeptide parvalbumin (PV) (**Fig. 3 A**). We have previously shown that these Cre lines direct RiboTag expression in the desired cell types (Kaczmarczyk et al., 2022). Since the PV Cre line targets the same cerebellar cell types as the Gad2 line, and were thus redundant, the cerebellar PV samples were not processed.

**Figure 3:**
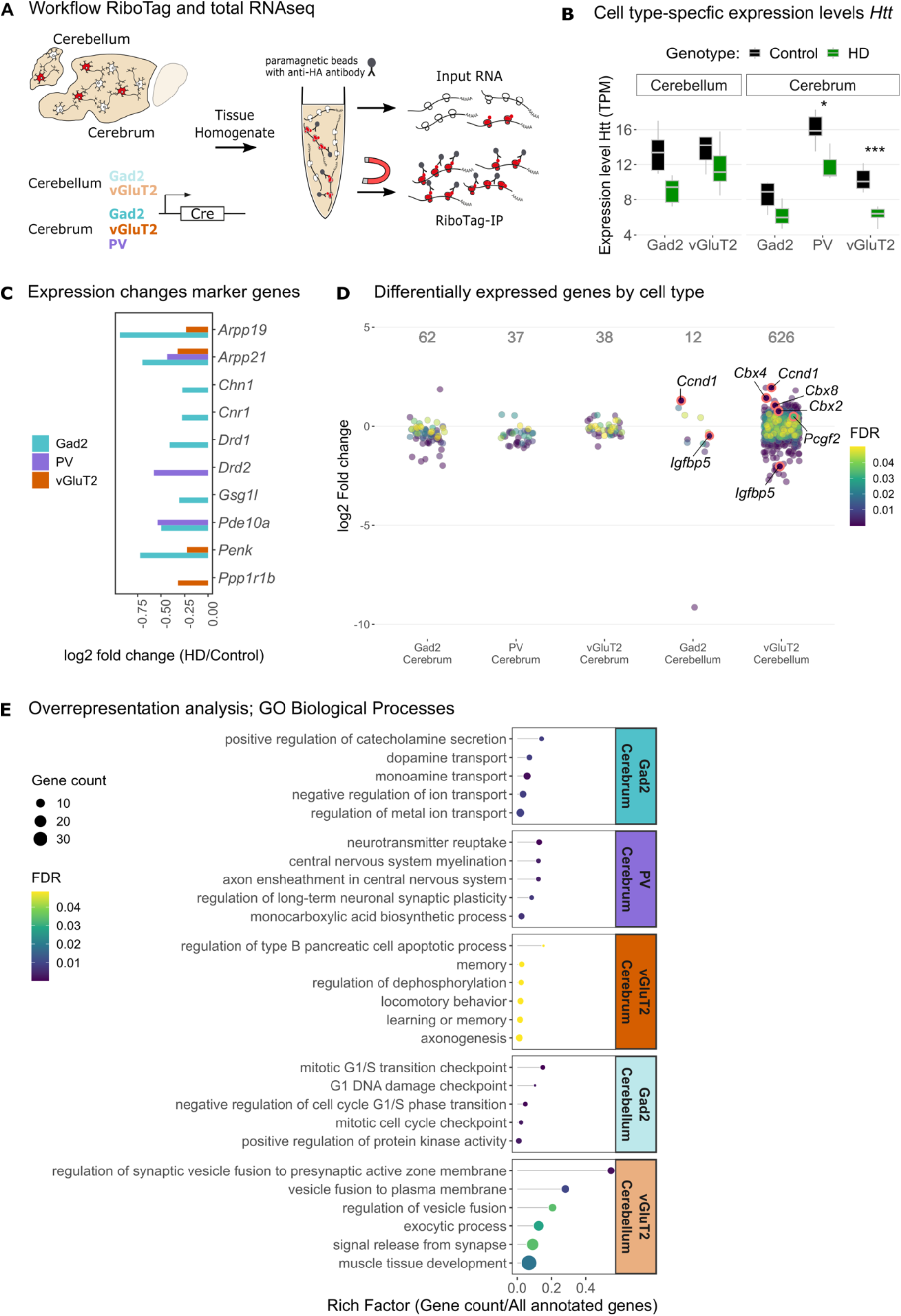
Translatome analysis with RiboTag reveals cell type specific responses in HD. **(A)** Hdh^Q7/Q200^ mice homozygous for RiboTag were crossed with homozygous Cre-driver lines to facilitate cell type-specific expression of Rpl22-HA in glutamatergic (vGluT2^+^) and GABAergic (Gad2^+^) neurons in the cerebellum and cerebrum, and parvalbumin (PV^+^)-expressing neurons in the cerebrum. Cerebellum and cerebrum (without olfactory bulb) were separated and flash frozen at 9 months. For RiboTag IPs, cell type-specific HA-tagged ribosomes were immunoprecipitated from tissue homogenate using anti-HA antibody bound to magnetic beads. Total RNA was prepared from an aliquot of the same tissue homogenate. **(B)** Expression levels of Htt mRNA were slightly reduced in all cell types, significantly in cerebral vGluT2^+^ and PV^+^ neurons (* FDR ≤ 0.05; ** FDR ≤ 0.01; *** FDR ≤ 0.001). **(C)** HD-typical genes showed downregulation in cerebral cell types. **(D)** Differentially expressed genes (DEGs) (FDR ≤ 0.05) for each cell type. **(E)** Top five significantly enriched terms (FDR ≤ 0.05) among DEGs reveals cell type-specific responses to mHtt.

Upon sequencing RiboTag-captured mRNAs, we found *Htt* mRNA levels were significantly reduced in cerebral PV^+^ and vGluT2^+^ neurons in HD mice, with the same trend in the other cell types (**Fig. 3 B**), consistent with the histological analysis (**Fig. 2 F**). Principal component analysis (PCA) of IP and total RNA samples showed a clear separation of RiboTag IP samples by targeted cell type, in contrast to total RNA samples (**Fig. S4 A, B**), indicating that we successfully obtained cell type-specific translatomes from RiboTag IPs. This was confirmed by analyzing expression of cell type-specific marker genes in comparison to total RNA samples (**Fig. S4 C, D**). General GABAergic markers *Gad2, Slc32a1, Dlx2, and Lhx6* were enriched in cerebral Gad2^+^ and PV^+^, but not vGluT2^+^ neurons (**Fig. S4 C**). PV^+^ samples showed enrichment for genes important in development and function of cortical PV^+^ interneurons (*Syt2, Cplx1, Nek7, Ank1, Kcnc2, Gpr176, Pthlh*) (Lucas et al., 2014; Hinojosa et al., 2018; Huntley et al., 2020; Stevens et al., 2021), while marker genes for Gad2^+^/PV^-^ neurons, including genes with high striatal expression levels, such as *Adora2a, Drd1, Drd2, Penk*, and *Ppp1r1b*, were depleted in PV^+^ samples, as were markers of other PV^-^ GABAergic interneuron subtypes (*Sst, Vip* and *Htr3a*). Cerebellar Gad2^+^ samples (**Fig. S4 D**) showed enrichment of markers for Purkinje (*Pcp, Grid2, Esrrb, Foxp2*), Basket (*Kcna2*), Golgi (*Lgi2, Gdj2, Sorcs3*) and Stellate cells (*Grik3*), while cerebellar vGluT2^+^ samples showed enrichment for granule cell markers (*Gabra6, Cntn2, Zic1, Etv1, Nfia*) (Kozareva et al., 2021; Tam et al., 2021). Astrocyte and microglia markers (Saudou and Humbert, 2016), were depleted in all IP samples compared to total RNA, as expected. Taken together, these results confirm successful isolation of cell type-specific translatomes with RiboTag, and inclusion of expected neuronal populations in both targeted regions.

### Translatome analysis reveals cell type-specific responses to mHtt

Next, we performed differential gene expression analysis to study the cell type-specific responses of targeted neurons to mHtt. For cerebral neurons, we observed downregulation of known HD-associated neuronal genes, such *as Drd1, Drd2, Penk, Ppp1r1b, Pde10a, Arpp21*, and *Pcp4, all* commonly reported to be downregulated in both human HD patients and in various HD mouse models (Gallardo-Orihuela et al., 2019; Malla et al., 2021; Świtońska-Kurkowska et al., 2021) (**Fig. 3 C**). Several of these genes show high striatal expression in early vulnerable populations i.e., Gad2^+^ SPNs, so that observed downregulation of these genes in cerebral Gad2^+^ neurons further support a striatal phenotype in our HD mice at 9 months. Comparison of our cell type-specific data with bulk RNAseq data from striata from a different HD KI model, zQ175DN (Langfelder et al., 2016), showed highly significant overlap between gene expression changes detected in cerebral Gad2^+^ samples and bulk striatal tissue at all observed time points (**Fig. S5 A**), indicating that our S4-HdhQ200 mice have HD-typical phenotypes. There was also significant overlap of bulk RNAseq data with our cell type-specific results for cerebellar Gad2^+^ and vGluT2^+^ neurons (**Fig. S5 B**). Notably, these HD-associated genes were also downregulated in cerebral vGluT2^+^ or PV^+^ neurons, which may be explained in at least two ways. First, downregulation of striatal marker genes may occur not only in highly vulnerable striatal projection neurons but in other cell types as well, as shown in a recent mouse study for glutamatergic corticostriatal projection neurons, astroglia, and cholinergic interneurons (Lee et al., 2020). Second, downregulation of these genes may occur in regions beyond the striatum, which would be detected when multiple regions are combined.

Surprisingly, cerebellar vGluT2^+^ neurons showed the overall highest number of differentially expressed genes (DEGs) with 626 (**Fig. 3 D**). To discover functional associations, we analyzed overrepresentation of gene ontology (GO) terms in the biological process classes among cell type-specific DEGs, which revealed cell type-specific responses (**Fig. 3 E**). DEGs of cerebral Gad2^+^ neurons were associated with dopamine transport and secretion, ion transport, downregulation of neuropeptides vasopressin and proenkephalin (*Vip, Penk*), and downregulation of several genes encoding neurotransmitter receptor components for dopamine (*Drd1*), glutamate (*Grm3*), cholinergic (*Chrna6*), cannabinoid (*Cnr1*), and serotonin (*Htr4*) (**Fig. S6 A**). Cerebral vGluT2^+^ neurons were enriched for gene sets associated with cognition, memory, and locomotory behavior, while PV^+^ neurons were associated with myelination and neurotransmitter uptake. Top enriched pathways were related to exocytosis and vesicle fusion (**Fig. 3 E**), driven by downregulation of genes encoding the central components involved in calcium-dependent synaptic vesicle fusion and exocytosis, such as complexin-encoding genes *Cplx2* and *Cplx3, Snap25*, and *Vamp2*, as well as differential expression of Ca^2+^ sensors *Syt7, Doc2b* and *Otof (***Fig. S6 A**). Cerebellar Gad2^+^ neurons, in contrast, indicated activation of mitotic cell cycle regulation, in particular G1/S transition, with upregulation of cyclin D1 (*Ccnd1*), a regulator of G1/S progression, downregulation of antiproliferative p21/cyclin-dependent kinase inhibitor 1a (*Cdkn1a*) and polo-like kinase 5 (*Plk5*) (**Fig. S6 A**). *Ccnd1* was also among the strongest upregulated genes in cerebellar vGluT2^+^ neurons (log2FC = 1.8, FDR < 0.001). This was validated by ISH for *Ccnd1* mRNA in tissue sections of 9 months old HdhQ200 mice **(Fig. 4 A)**, showing a 1.7-fold increase in *Ccnd1* staining in the granule layer in HD mice (p = 0.008) (**Fig. 4 B**). Therefore, the cerebellum shows a robust, cell type-specific response to the HD mutation as early as 9 months with a focus on pathways involving neurotransmission or cell cycle reentry.

**Figure 4:**
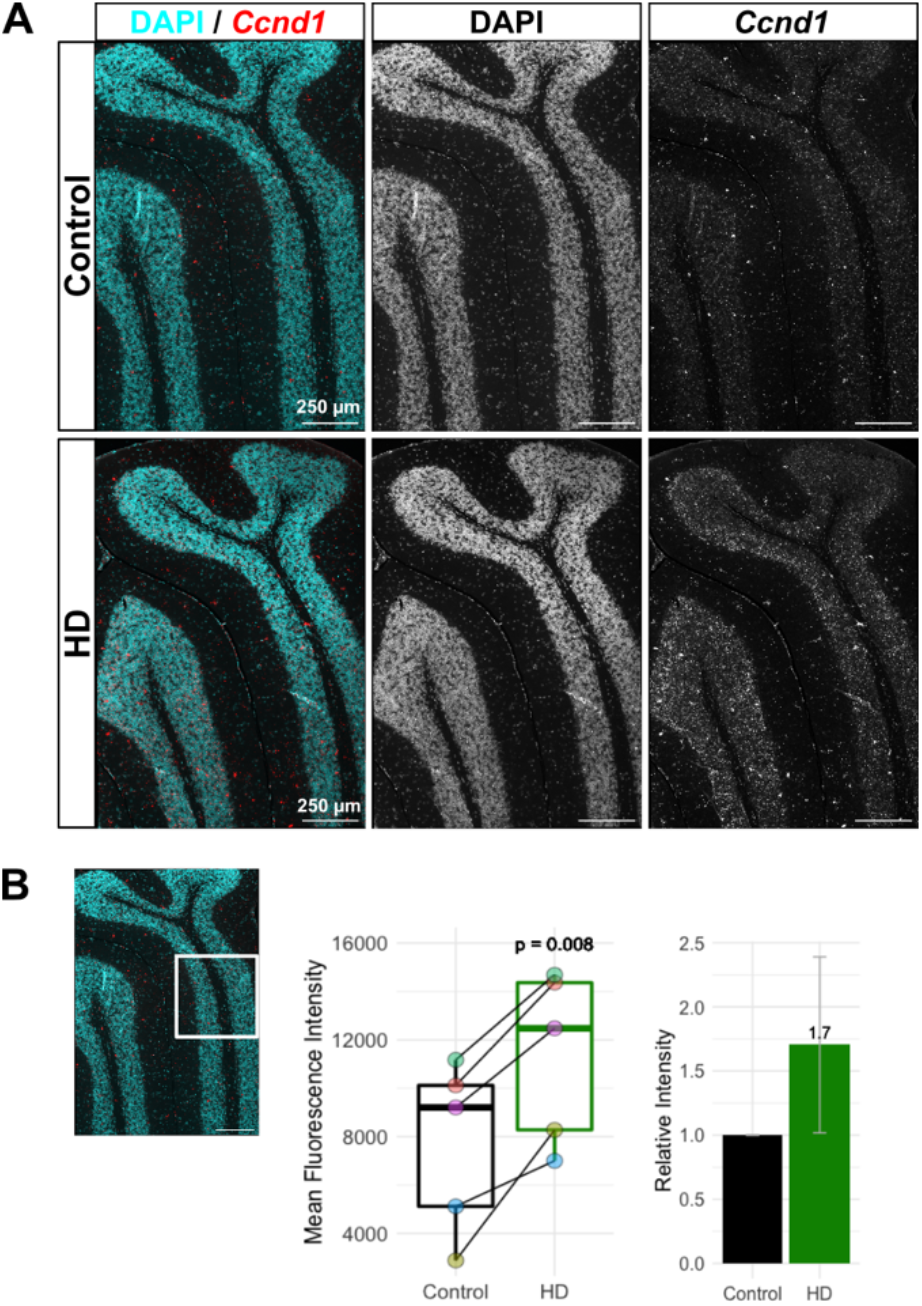
Ccnd1 ISH staining is increased in the granular layer of 9-month-old HD mice. *Ccnd1* mRNA signal is increased in HD cerebella at 9 months. (A) *In situ* hybridization for *Ccnd1* mRNA (red) shows increased signal in the anterior cerebellar lobules III-V in 9-month-old HD mice (bottom) compared to littermate controls (top). (B) Quantification of mean fluorescence intensity in the granule layer of lobule IV/V (white square) from matched HD and Control at 9 months shows a significant 1.7-fold increase of *Ccnd1* staining in HD mice (p = 0.008 paired t-test, n = 5, matched Control and HD samples are indicated by dot color and connecting line)

### Cerebellar neurons show cell type-specific responses to mHtt

Given the high number of DEGs detected in cerebellar vGluT2^+^ neurons and that nearly all of the cerebellar Gad2^+^ DEGs were also changed in vGluT2^+^ cerebella, we conducted additional analyses of expression changes. Comparison of DEGs between cerebellar cell types revealed that the shared DEGs changed with the same directionality (**Fig. S6 B**), again suggesting a similar response. We next performed gene set enrichment analysis (GSEA) on both cell types, using six different statistical methods to calculate enrichment and a consensus score which was used to rank results (Väremo et al., 2013). GSEA is a useful tool to detect pathways that have a coordinated response even when individual genes of that pathway have small and statistically insignificant changes. Interestingly, the GSEA indicated that, despite sharing the majority of the DEGs with cerebellar vGluT2^+^ neurons, cerebellar Gad2^+^ neurons had a remarkably different response (**Fig. 5**). Genes associated with “translation and co-translational targeting of proteins to membrane” and “endoplasmic reticulum” were enriched in both cerebellar cell types (**Fig. S7**). However, Gad2^+^ neurons uniquely showed upregulation of protein glycosylation-related gene sets and ATP synthesis, oxidative phosphorylation, and Parkinson’s disease pathway, suggesting an increase in mitochondrial function in these neurons. In contrast, vGluT2^+^ cerebellar neurons made a cell type-specific response in the form of upregulation of cell differentiation-associated gene sets, cell cycle regulation and chromosome organization, autophagy, metabolic processes, upregulation of PI3K-Akt signaling pathway genes and apoptosis. This suggests that, despite superficial similarities, cerebellar cell types activate different pathways in response to mHtt. Additionally, the enrichment of genes associated with differentiation, PI3K-Akt signaling, and apoptosis, together with the upregulation of *Ccnd1*, indicate that cell cycle regulation is affected in cerebellar vGluT2^+^ neurons. Interestingly, we found significant overlaps of DEGs identified in both vGluT2^+^ and Gad2^+^ cerebellar neurons with DEGs identified in the cerebellum of a knock-in mouse model of Spinocerebellar ataxia 1 (SCA1) [OMIM:164400], an autosomal dominant disease caused by expansion of a polyglutamine encoding CAG repeat, like in HD, but in the Ataxin1 gene (*Atxn1*) (**Fig S8**) (Driessen et al., 2018). Genes sharing expression patterns in both vGluT2^+^ and Gad2^+^ cerebellar cell types were also among the earliest DEGs in SCA1 (**Fig S8**).

**Figure 5:**
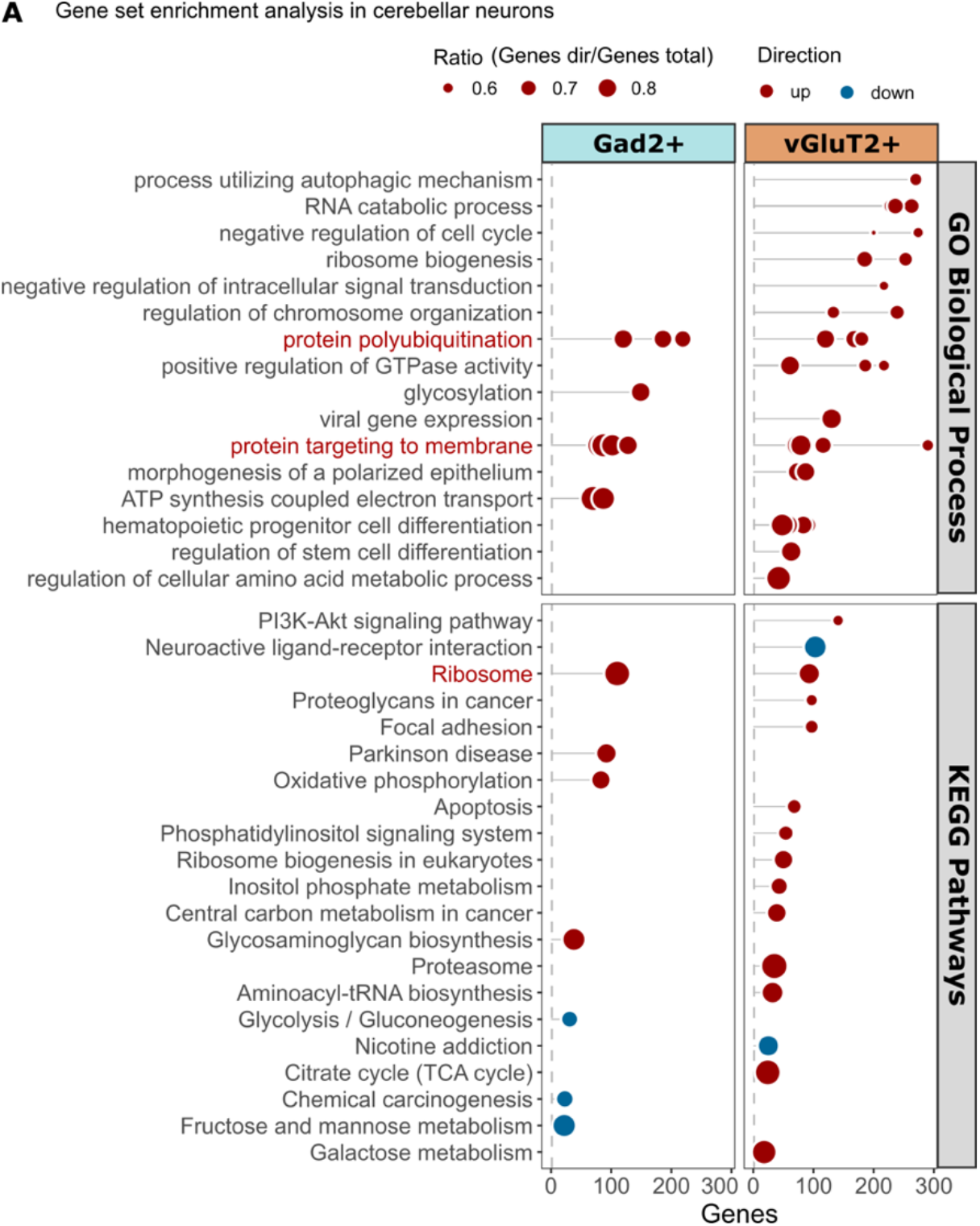
Cerebellar neurons show early and cell type-specific responses. Gene set enrichment analysis shows a largely cell type-specific response in Gad2^+^ and vGlut2^+^ cerebellar neurons, shared terms are indicated in red. GO terms were reduced to parent terms based on semantic similarity. Ratio indicates the relative number of enriched genes to gene set size.

### Polycomb repressor complex 1 (PRC1) protein genes are upregulated in cerebellar vGluT2+ neurons

We next investigated if the differential expression patterns we observed in cerebellar vGluT2^+^ neurons could be driven by specific transcription factors using the ChIP Enrichment Analysis (ChEA) database (Lachmann et al., 2010) to perform overrepresentation analysis against the transcription factor targets determined by Chip-X. This analysis revealed that 259 out of 626 DEGs were among target genes of polycomb repressor complex 2 (PRC2) components SUZ12 or EZH2, PRC1 core component RING2B/RNF2, or PRC-associated factor MTF2 (**Fig. 6 A**). Of these 259 genes, 218 were associated with PRC2 core proteins EZH2 (97 DEGs) and SUZ12 (207 DEGs), and 131 DEGs with PRC1 protein RING2B/RNF2 showing near equal distribution of up- and downregulated DEGs (**Fig. 6 B**). Levels of PRC2 core components and associated factors were not affected, but we found increased expression of canonical PRC1 complex components chromobox proteins *Cbx2, Cbx4, Cbx8* and polycomb group ring finger 2 (*Pcgf2)* in cerebellar vGluT2^+^ neurons (**Fig. 6 C**). Although PRC2 regulation has been shown to be impaired in HD (Seong et al., 2009; Francelle et al., 2017), little is known on the involvement of PRC1 in HD. However, given the proposed roles of CBX proteins in regulating cell cycle progression across various checkpoints (van Wijnen et al., 2021), these findings further support the view of aberrant cell cycle regulation in cerebellar vGluT2^+^ neurons.

**Figure 6:**
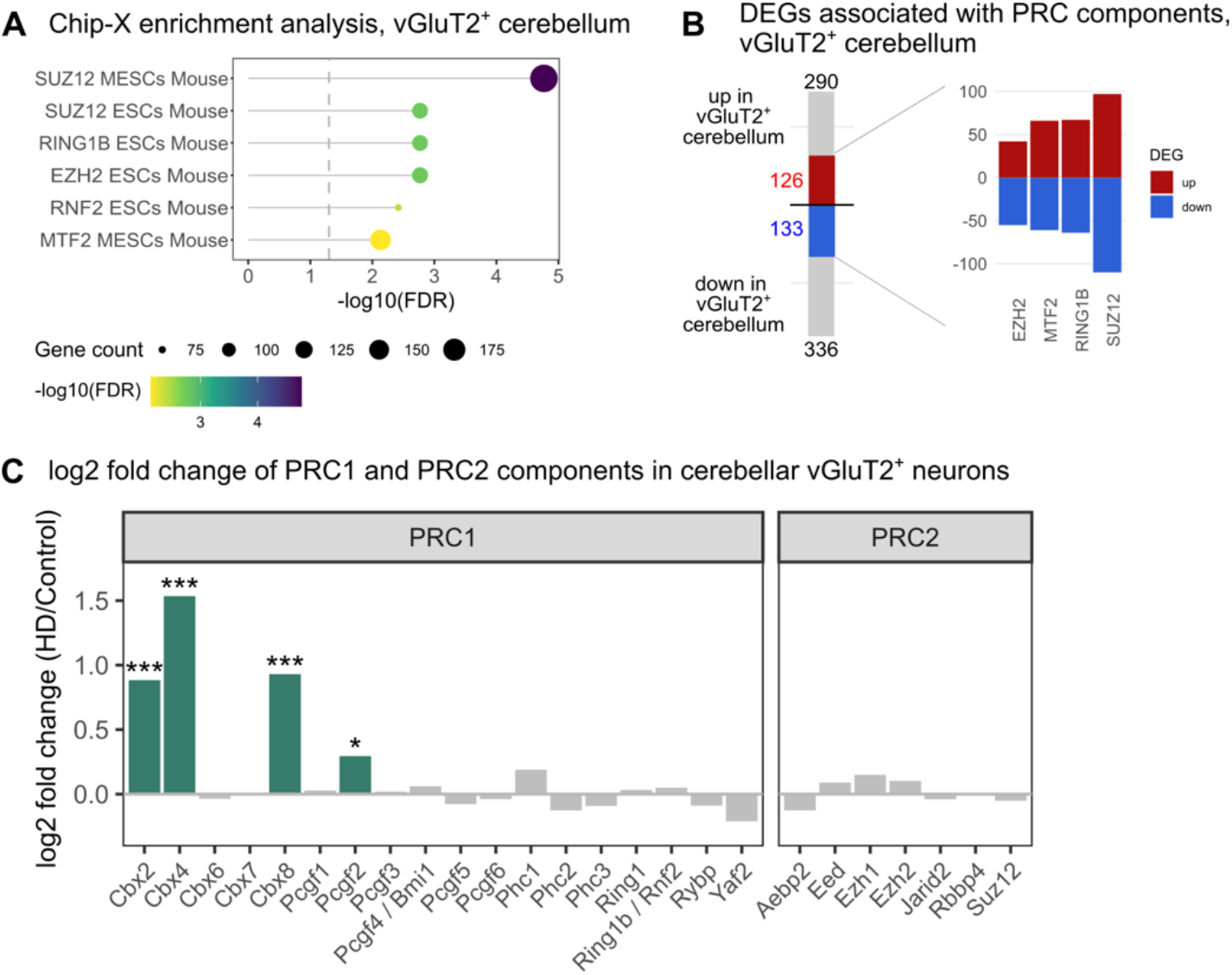
Polycomb repressor complex 1 (PRC1) proteins are upregulated in cerebellar vGluT2+ neurons. **(A)** Overrepresentation analysis against ChEA ChIPseq gene sets from mouse embryonic stem cells (ESCs/MESCs) show DEGs detected in vGluT2^+^ neurons are overrepresented among targets associated with polycomb repressor complex (PRC) proteins (FDR ≤ 0.01). **(B)** PRC-associated DEGs show similar distribution between up and down-regulated DEGs. **(C)** PRC1 protein genes *Cbx2, Cbx4, Cbx8* and *Pcgf2* are significantly upregulated in cerebellar vGluT2^+^ neurons (green) in HD, whereas PRC2 core protein genes (*Eed, Ezh1, Ezh2, Suz12*) and associated factors (*Aebp2, Jarid2, Rbbp4*) are not differentially expressed. (* FDR ≤ 0.05; ** FDR ≤ 0.01; *** FDR ≤ 0.001)

## DISCUSSION

Here we characterized an HdhQ200 knock-in mouse model in a new background, 129S4. Behavioral and neuropathological characterization of this mouse model showed widespread Htt^+^ NIIs and downregulation of *Htt* and *Ppp1r1b* mRNAs by 9 months, with a very mild behavioral phenotype at later disease stages. These histological changes motivated us to use RiboTag to further analyze 9 months old HD mice, and to identify, for the first time, cell type-specific translatome responses in the cerebellum at an early disease stage.

Surprisingly, our translatome analysis revealed widespread changes in cerebellar vGluT2^+^ neurons, a population consisting predominantly of granule neurons. This result was unexpected, as granule cells are typically considered resistant in HD. Instead, compromised function and loss of Purkinje cells is the cerebellar phenotype most described in HD patients and mouse models (Jeste et al., 1984; Dougherty et al., 2013; Rüb et al., 2013). Functional analysis of DEGs in cerebellar vGluT2^+^ neurons revealed vesicular fusion and exocytosis as predominantly enriched biological processes, suggesting neurotransmitter release and synaptic signaling may be affected. GSEA further suggested upregulation of genes related to differentiation, cell cycle regulation, and energy metabolism.

One of the most notable gene expression changes was the strong upregulation of *Ccnd1* in both cerebellar vGluT2^+^ and Gad2^+^ neurons, which was further confirmed by ISH staining. Ccnd1 is a core regulator of G1/S transition, and its upregulation is associated with cell cycle reentry (CCR). CCR of postmitotic neurons has been proposed as an early event in progressive neurodegeneration which can be triggered by dysregulation of various mechanisms, several of which are known to be affected in HD, e.g., altered levels of neurotrophic factors, oxidative stress, unrepaired DNA damage, or dysfunction of the ubiquitin-proteasome system (Joseph et al., 2020). Activation of Notch signaling and the downstream Akt/GSK3b pathway in response to excitotoxic stimulus can cause CCR in mature neurons by Ccnd1 upregulation driving G1/S transition, which can in turn prompt neurons to enter apoptosis (Marathe et al., 2015). Indeed, Ccnd1 upregulation has also been observed in striatal neurons in response to 3-Nitropropionic acid treatment (Pelegrí et al., 2008; Dietrich et al., 2022), a model replicating HD-like striatal neuron loss. This explanation is in line with our observation of upregulation of PI3K-Akt signaling and apoptosis-related genes in cerebellar vGluT2^+^ neurons, suggesting CCR may also occur in cerebellar vGluT2^+^ neurons, at early, pre-symptomatic disease stages.

Another important finding was the high percentage of DEGs in cerebellar vGluT2^+^ neurons associated with PRC components. HTT is a known facilitator of PRC2 (Seong et al., 2009; Biagioli et al., 2015; Dong et al., 2015) and neurons expressing mHtt show aberrant chromatin landscapes already at early developmental stages, which may result in reduced neuronal fitness (Biagioli et al., 2015), and PRC2 silencing is important for maintaining cell identity in differentiated neurons (Von Schimmelmann et al., 2016; Malaiya et al., 2021). Given the established role of PRC2 in HD, the high number of PRC2-associated DEGs in cerebellar vGluT2^+^ neurons suggests an epigenetic mechanism underlies the observed changes. In contrast, little is known on whether PRC1 plays a role in HD. Therefore, we were surprised to see upregulation of several canonical PRC1-related genes in cerebellar vGluT2^+^ neurons. Cbx family proteins (Cbx2,4,6,7,8) form core elements of the canonical PRC1 complex recognizing H3K9me2 marks. Beyond this, they control cell cycle, proliferation, and senescence with Cbx2, Cbx4 and Cbx8 having distinct roles in regulating cell cycle progression (van Wijnen et al., 2021). Cbx2 and Cbx8 may further act as positive regulators of axonal regeneration (Duan et al., 2018), while Cbx4, a SUMO E3-ligase, is essential for SUMOylation of Bmi1/Pcgf4, facilitating its recruitment and accumulation at DNA damage sites (Ismail et al., 2012). Additionally, Cbx2 and Cbx8 proteins also interact with REST, a master regulator of neuronal gene expression and interaction partner of Htt, affecting PRC1 binding patterns and REST-mediated transcriptional repression (Ren and Kerppola, 2011). This functional diversity of CBX proteins and their involvement in central mechanisms that are disturbed in HD, such as cell cycle regulation and DNA damage repair, provides several interesting mechanisms by which upregulation of these genes may be important in the response of granule cells, opening new avenues for future research.

Interestingly, our results from specific cerebellar neurons of HD mice also had considerable overlap with bulk RNAseq data from cerebella of a SCA1 mouse model (Driessen et al., 2018). Most notably, the pronounced downregulation of *Igfbp5* in the cerebellum was also described as a robust feature in SCA1 and SCA7 knock-in mice (Gatchel et al., 2008). Igfbp5 downregulation increases IGF1 availability, which may trigger pro-survival pathways via IGF1R signaling to support neuronal function (Roschier et al., 2001; Gatchel et al., 2008). This suggests mechanistic similarity between the polyglutamine diseases HD and SCA1, a finding that would be interesting in the light of an increased understanding of cerebellar involvement in HD pathology and symptoms in various HD patients.

Further research is required to determine the mechanism and consequences of the translatome changes observed in our experiments, especially regarding the question whether the observed responses are detrimental to neurons or an adaptive response. However, the pronounced cell type-specific response in cerebellar vGluT2^+^ neurons early in disease offers new and interesting insights into early disease-response in the cerebellum. While the presence and extent of cerebellar dysfunction in HD appears to be more variable than striatal perturbations, it also opens the possibility of finding new potential therapeutic targets and interventional strategies. Indication of cerebellar compensation in early stages of HD make this approach particularly appealing.

Finally, one of the goals of the study was to reveal new insight into the phenomenon of selective vulnerability. While we uncovered molecular pathways underpinning the cerebellum’s response to HD, the understanding of the broader topic of selective vulnerability was only modestly enhanced. Nonetheless, we found it interesting that the thalamus is the brain region with the second highest Htt expression but is a region with few Htt aggregates, while the striatum and cerebellum are low expressors but are highly prone to form aggregates, especially NIIs. These observations suggest that HD vulnerable brain regions are poorly equipped to handle the Htt protein in general, and the inclusion of a long polyglutamine stretch makes that struggle more severe.

### Limitations

One limitation of our experimental setup is that we studied ribosome-bound, translating mRNA and thus are not able to conclude whether observed changes occur due to changes at a transcriptional or translational level. This could be addressed by methods such as the Tagger mouse line (Kaczmarczyk et al., 2019), enabling cell type-specific assessment of four levels of gene expression. Another limitation is that we studied cell populations with various degrees of heterogeneity. This may partially explain the high number of DEGs in cerebellar vGluT2^+^ neurons, arguably the least heterogeneous cell type analyzed in this study. However, in a previous study using the same experimental approach, we did not observe strong changes in the cerebellar vGluT2^+^ neurons (Bauer et al., 2022), indicating this result is not an artifact but reflects the biological response of this cell type in HD. A third limitation is the use of a genetic background not widely used in HD research. In previous experiments, we observed that a subset of mice with a C57BL/6 background were hyperactive at night, which we thought could impact gene expression (Kaczmarczyk et al., 2021). To this end, the HdhQ200 model was backcrossed to a 129S4 background. This model has a very mild behavioral phenotype, like other knock-in HD mouse models. Importantly, histological assessment and translatome analysis revealed an HD-typical phenotype and many gene expression changes we detected have been detected in other studies of HD mice. Although direct comparison of our results from those of other studies require consideration of possible genetic background effects, the existence of the HdhQ200 model on a new genetic background may be useful for identifying genetic modifiers.

## METHODS

Ethical permissions for this work were granted by the Linköpings djurförsökestiska nämnd (permission # 14741-2019) and the Landesamt für Natur, Umwelt und Verbraucherschutz Nordrhein-Westfalen (permission #s 84-02.04.2013.A169 and 84-02.04.2013.A128).

In all these studies, HD mice were heterozygous as the very long mutation may partially inactivate the gene’s native function, and homozygotes may partially develop *Htt* knock-out phenotypes. Additional details are provided in **supplemental methods**.

### For RiboTag and histology experiments

129S4-HdhQ200 mice were crossed three times to homozygous RiboTag mice (B6N.129-Rpl22^tm1.1Psam^/J, line #011029; Jackson Laboratory, Bar Harbor, ME), on a 129S4 background (> 99.5% 129S4) to establish Hdh^Q200/Q7^/Rpl22-HA^flox/flox^ mice. Cre-driver lines vGluT2-IRES-Cre (Vong et al., 2011) (Slc17a6^tm2(cre)Lowl^/J, line #016963; Jackson Laboratory), Gad2-IRES-Cre (Haimon et al., 2018) (Gad2^tm2(cre)Zjh^/J, line #010802; Jackson Laboratory), and PV-IRES-Cre (Hippenmeyer et al., 2005) (B6;129P2-Pvalb^tm1(cre)Arbr^/J, line #008069; Jackson Laboratory), were each backcrossed to be > 99% 129S4 (Kaczmarczyk et al., 2021). To obtain experimental animals, male Hdh^Q7/Q200^/Rpl22-HA^flox/flox^ mice were crossed with female Hdh^Q7/Q7^ mice homozygous for Cre, producing offspring that are heterozygous for Cre and RiboTag, and either heterozygous or wild-type for *Htt*. Mice were sacrificed at approximately 9 months (average: 41.5 weeks, SD: 1.8; range: 39-45 weeks) by CO2 asphyxiation. The olfactory bulb was removed from the second hemisphere and cerebellum and cerebrum were separated by dissection, flash frozen on dry ice, and stored at -80 to -72 °C. Details are also provided in Supplementary Dataset 1. The most abundant CAG repeat lengths were measured for each sample with a fragment analyzer. The smallest, median, and longest repeat in the sample population: vGluT2 = 203, 205, 209; Gad2 = 201, 203, 207; PV = 202, 204, 207.

### For behavioral experiments

HdhQ200 mice were bred to 129S4 to generation 12 and subsequently bred to 129S4 mice carrying the Disrupted in schizophrenia 1 (*Disc1*) and non-agouti genes from C57Bl/6NTac mice that were 99.8% 129S4. The smallest, median, and longest repeat in the sample population: 160, 166, 174.

### Behavior

All tests were performed on the same cohort of control (3 male, 4 female) and HD mice (7 male, 5 female) between 3-18 months. Following 2 initial training days, motor tests were performed every 2-3 months and animals were weighed monthly. Mice were tested on the **accelerating rotarod** (ENV-574M, Mead Associates Inc, Fairfax, VT) for 300 seconds (s), with acceleration every 30 s from 4 revolutions per minute (rpm) to 40 rpm and latency to fall was recorded. Followed by the **balancing beam** paradigm, in which mice crossed an ascending, narrow beam (1 m, 17° ascent and 1.5 -0.5 cm taper across the width) to reach a black box at the high end and time needed to traverse the beam was recorded. At each time point, mice performed two repeat balancing tasks, interspersed by burrowing. **Burrowing behavior** of animals was tested for 1 h in light phase. Individual mice were placed in a cage with 200 ml of standard bedding substrate and a burrow made of a 20 cm long black plastic tube with a diameter of 7 cm, with the open end raised 3 cm and the closed end raised 1 cm from the cage floor. Removed bedding material was used as a measure of burrowing activity.

### Neuropathology

#### Tissue preparation

Mice were sacrificed by CO^2^ asphyxiation followed by transcardial perfusion with 10% formalin solution. Dissected brains were separated into hemispheres along the midline and fixed for 2 days at 4°C with gentle shaking. Fixed brains were paraffinized with each cassette containing age-matched HD and control samples and cut into 4 μm sections.

#### Immunohistochemistry/Immunofluorescence staining

Tissue sections were deparaffinized and boiled in citrate buffer (pH 8) for 20 min for antigen retrieval, then cooled for 20 min at room temperature (RT). For NII staining, sections were incubated with 25 μg/ml proteinase K at 30 min/37°C to remove cytosolic HTT. Sections were treated with 0.3% H^2^O^2^ for 30 min/RT, then permeabilized with blocking buffer (PBS with 2.5% normal horse serum (NHs)) for 1 h/RT, followed by incubation with primary antibody for 30 min/RT. Sections were washed twice in PBS for 5 min and secondary antibody was incubated for 30 min/RT, followed by 2x 5 min PBS washes. For immunohistochemistry, sections were colorized using NovaRED (SK-4800, Vector Laboratories, Burlingame, CA). Primary antibodies: Huntingtin 1:100 (ab109115, Abcam, Cambridge, UK); GFAP 1:500 (GA524, Dako Omnis); Iba1 1:200 (019-10741, Wako Chemicals). Fluorescent secondary antibodies: Alexa Fluor 488 donkey anti-rabbit 1:250 (Jackson ImmunoReseach, AB_2313584).

***In situ* hybridization (ISH)** was performed according to the RNAScope Fast-RED protocol followed by autofluorescence quenching for 5 min with TrueView (Vector Laboratories). Counterstaining with DAPI (1:20000 in PBS with 2.5% NHS) was performed to analyze co-localization of *Htt*.

#### Image analysis and quantification

For *Htt* ISH, images of sections were taken with a Zeiss Axio A1/D1 microscope (Zeiss, Oberkochen, Germany) and deconvoluted using Huygens software (Scientific Volume Imaging (SVI), Hilversum, Netherlands). For quantitative analysis of Htt aggregates, *Htt* RNA staining, and HTT/DAPI colocalization, particle counting was done using IMARIS with default settings (Bitplane, Oxford Instrument, England). For *Ppp1r1b* ISH integrated density was measure in FIJI. *Ccnd1* ISH quantification was performed on matched HD and Control sections, from the same cassette and imaged using a Zeiss LSM700 confocal microscope with identical settings. For quantification, a sum projection of Z-stacks was performed in FIJI and mean fluorescence intensity was measured in the granular layer after background correction and averaged for each section.

### RiboTag RNAseq

#### Preparation of RNA samples

For RiboTag samples, cell type-specific mRNA was immunoprecipitated from brain tissue homogenates of flash-frozen cerebellum or cerebrum hemispheres, as described here (Bauer et al., 2022). In brief, homogenate was precleared by centrifugation and incubation with IgG isotype antibody-bound protein-G dynabeads (PGDB; 1009D, Invitrogen, Waltham MA). HA-tagged ribosomes with bound mRNAs were captured by incubation with 12CA5 anti-HA antibody (11666606001, Roche) 90 min/4°C, followed by incubation with PGDBs for 60 min/4°C. Beads were washed using buffers with increasing salt concentration and mRNA was eluted by phenol-chloroform extraction, followed by clean up using the Qiagen RNeasy Mini kit. Prior to performing the RiboTag protocol, total RNA was isolated from tissue homogenates as input control.

#### Sequencing and alignment

Illumina TruSeq Stranded mRNA libraries were sequenced (paired end, 100 bp) on a NovaSeq6000 (Illumina, San Diego CA) on a S4 flow cell. Library preparation and sequencing was performed at NGI Uppsala, Sweden. Alignment and mapping was performed with STAR/Salmon. Samples were kept if they contained >24M uniquely mapped reads.

#### Bioinformatic analysis

**For principal component analysis**, the variance for protein-coding genes was calculated based on log-scaled transcripts per million (TPM) values across RiboTag IP or total RNA samples. Top 500 most variable genes were used for PCA. To analyze enrichment of cell type-specfic marker genes in RiboTag IP samples, we normalized log2-transformed TPM values to total RNA and calculated the row-wise z-score: Z = (x – mean(total RNA))/SD(row), where x is the sample value and SD is the row-wise standard deviation. Heatmaps were visualized with pheatmap (v1.0.12).

**Differential expression analysis** was performed with DESeq2 (v1.30.1) comparing disease and control samples for each cell type. Only protein coding genes with a row-wise mean count > 10 were considered. Genes with a false discovery rate (FDR)-adjusted p-value ≤ 0.05 were considered as differentially expressed.

**Overrepresentation analysis (ORA)** for DEGs was performed for each cell type using clusterProfiler (v3.18.1), with a cutoff of FDR < 0.05.

**Gene set enrichment analysis (GSEA)** for GO Biological Process (c5.go.bp.v7.4.symbols; gsea-msigdb.org) and KEGG pathways (KEGG_mouse_2019; maayanlab.cloud/Enrichr) was performed with piano (v2.6.0) (Väremo et al., 2013) using six methods (“mean”, “median”, “sum”, “stouffer”, “reporter” or “tailStrength”) to calculate statistical significance. Median consensus scores were calculated based on adjusted p-values using the integrated consensusScores function. Terms with distinct directional FDR ≤ 0.05 in at least two of the six applied gene set statistics were included. For visualization, GO terms were collapsed to parent terms by semantic similarity (Resnik, threshold = 0.8) using rrvigo (v1.2.0).

#### Chip-X enrichment analysis (ChEA)

(Lachmann et al., 2010) was performed using the enrichr R-package (v3.0) to analyze overrepresentation of DEGs among gene sets in the ChEA 2016 collection of transcription factor-associated genes.

**Overlap analysis** of cell type-specific DEGs from RiboTag IPs with bulk RNAseq data was calculated using Fisher’s exact test with the GeneOverlap R package (v1.26.0), using the average number of protein coding genes detected in IP samples (12519) as background. External data sets: HD [GEO:GSE65776]; SCA1 [GEO:GSE122099].

## DATA ACCESS

All raw and processed sequencing data generated in this study have been submitted to the NCBI Gene Expression Omnibus (GEO; https://www.ncbi.nlm.nih.gov/geo/) under accession number GSE199837.

## ACKNOWLEDGMENTS

The authors would like to thank the National Genomics Infrastructure (NGI) Uppsala, Sweden for aiding with library preparation and sequencing. Computations and data handling were enabled by resources provided by the Swedish National Infrastructure for Computing (SNIC) at UPPMAX, partially funded by the Swedish Research Council (2018-05973). We would like to thank Rui Benfeitas and National Bioinformatics Infrastructure Sweden (NBIS) for support provided through the Swedish Bioinformatics Advisory program. We also thank David Engblom for access to equipment, Maria Ntzouni for preparing histological samples, and Åsa Schippert and Mouna Tababi for their help with repeat sizing. This work was supported by the German Center for Neurodegenerative Diseases, the Wallenberg Center for Molecular Medicine, and the Knut and Alice Wallenberg Foundation (KAW 2019-0047).

## AUTHORS CONTRIBUTION

MJ performed behavioral experiments and characterization. C-YC performed histopathological characterization and analysis. SM performed repeat sizing. LK collected tissue samples. SB performed RNA sample preparation, bioinformatic analysis and ISH. WSJ devised the model and designed and supervised the study. The manuscript was written by SB and WSJ with input from all authors.

